# Tribbles 2 confers enzalutamide resistance in prostate cancer by promoting lineage plasticity

**DOI:** 10.1101/2021.03.26.437250

**Authors:** Jitender Monga, Indra Adrianto, Craig Rogers, Shirish Gadgeel, Dhananjay Chitale, Joshi J. Alumkal, Himisha Beltran, Amina Zoubeidi, Jagadananda Ghosh

## Abstract

Second-generation anti-androgen, such as enzalutamide (Xtandi), is commonly prescribed for prostate cancer therapy, but enzalutamide-resistant, lethally incurable disease invariably develops. To understand the molecular basis of enzalutamide resistance, we comprehensively analyzed prostate tumors and clinically relevant models. These studies revealed that enzalutamide resistant prostate cancer cells overexpress Tribbles 2 (Trib2), a pseudokinase. Expression of Trib2 is negatively regulated by androgen receptor signaling. Overexpression of Trib2 makes prostate cancer cells completely resistant to clinically relevant doses of enzalutamide. Trib2 downregulates expression of luminal markers but upregulates the neuronal transcription factor, BRN2, and the stemness factor, SOX2, to induce neuroendocrine differentiation. Our findings indicate that Trib2 confers resistance to enzalutamide therapy via a mechanism involving increased cellular plasticity and lineage switching.

## Main Text

Common forms of prostate cancer cells bear luminal characteristics and depend on androgenic signaling for growth. However, it has been realized that men with prostate cancer who were treated with anti-androgenic therapies, frequently develop aggressive and deadly forms of prostate cancer which are no longer responsive to primary therapies. Enzalutamide, an inhibitor of androgen receptor function, is commonly prescribed to treat prostate cancer, but resistant prostate cancer eventually develops which grow aggressively, leading to widespread metastatic disease and bring demise to prostate cancer patients (*1–3*). Based on current assessment, the enzalutamide-resistant type of aggressive prostate cancer is responsible for most of the morbidity and mortality associated with prostate cancer and ~30,000 lives of American men are lost every year (*4*). However, lack of proper understanding about critical molecular targets in hormone refractory disease settings, such as enzalutamide resistance, largely contributes to the loss of battle against majority of prostate cancer deaths.

Several reports encompassing the involvement of both androgen receptor reactivation or bypass, as well as androgen-independent signaling, have been forwarded to explain the mechanism of enzalutamide resistance. However, analysis of multiple cell lines and *in vivo* models, which were used to explore the molecular basis, have ended up with identification of subtypes of cells (*5–7*). Current molecular understanding suggests that in addition to AR reactivation by mutation or splice variants, manifestation of enzalutamide resistance can be the result of overgrowth of cells that are developed in the tumor by lineage switching which is triggered by drug-induced repression or loss of the AR-signaling (*8–11*). About 10-20% of enzalutamide-resistant prostate cancer show neuroendocrine (NE) features and no effective treatments are available for this type of aggressive and highly invasive prostate cancer (*12–15*). Though the continued growth of the heavily enzalutamide-treated prostate tumors can be driven by non-androgenic signaling, molecular underpinnings of critical targetable mechanisms in treatment-emergent NE cells are yet to be fully characterized.

When prostate cancer cells become resistant to strong androgen receptor blocker, such as enzalutamide, their common characteristics change from slow-growing, non-invasive type to fast-growing, highly invasive cells, but identity of molecular drivers for their rapid growth and resistance to enzalutamide is still limited. To better understand the mechanistic basis behind enzalutamide resistance, we developed an *in vitro* model by chronically treating human LNCaP and MDA-PCA-2B prostate cancer cells (AR-signaling intact) with gradually increasing doses of enzalutamide (up to 30μM) for >12 weeks to mimic the clinical conditions in long-term enzalutamide therapy (*16*). Resultant cells (LNCaP-ENR and PCA-2B-ENR) are completely resistant to clinically relevant doses of enzalutamide, the blood level of which may go up to ~34μM in average (*17, 18*). An unbiased, comprehensive gene expression analysis revealed that the Trib2 pseudokinase is grossly overexpressed in enzalutamide-resistant prostate cancer cells, compared to parental enzalutamide-sensitive cells (Fig. 1a-d).

**Fig. 1.**
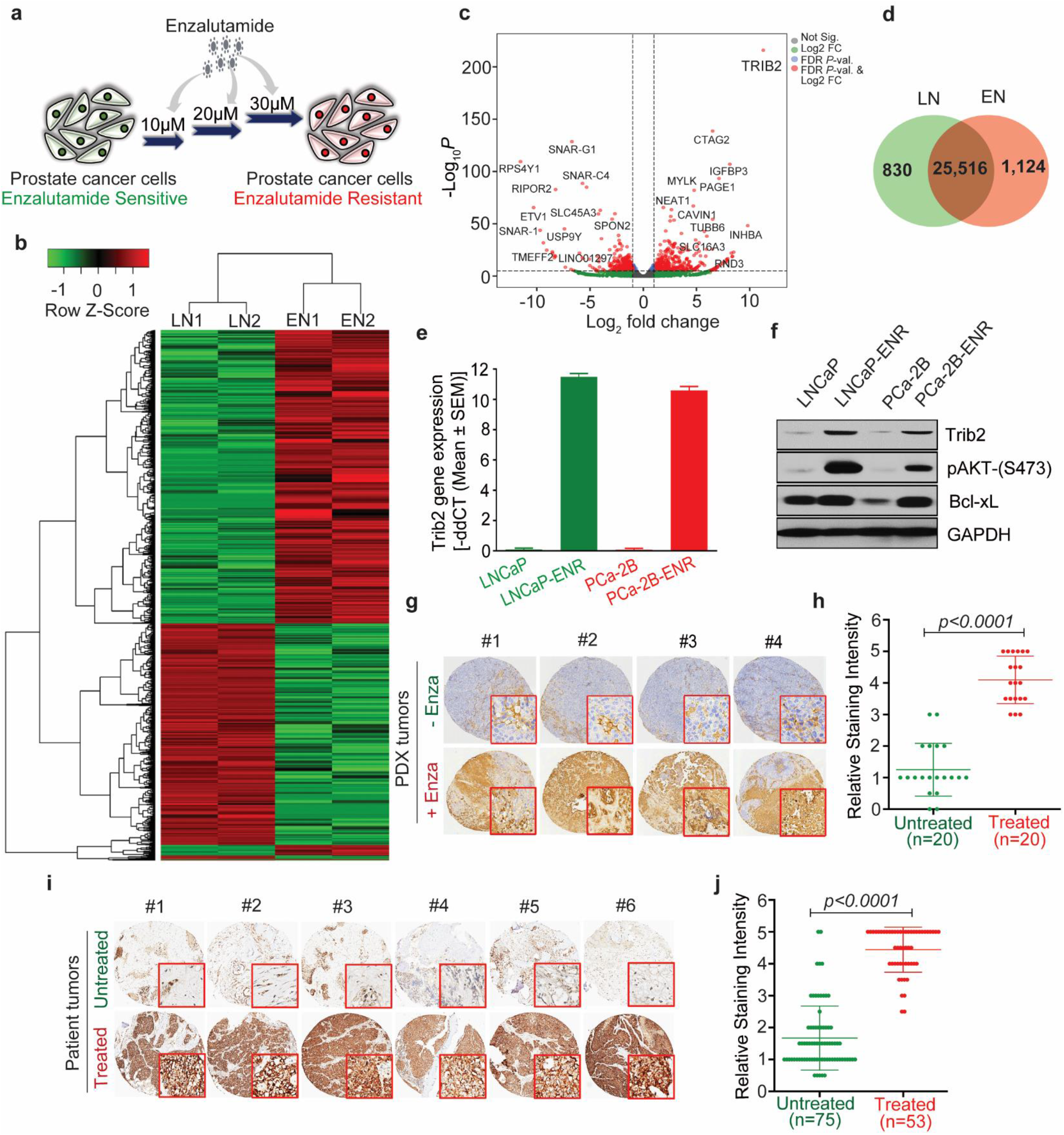
Overexpression of Trib2 in enzalutamide-resistant prostate cancer cells and tumors. **a** A model depicting the strategy to develop enzalutamide-resistant prostate cancer cells *in vitro*. **b** Heatmap showing up and downregulated genes in LNCaP-ENR cells (EN1 and EN2) compared to parental LNCaP cells (LN1 and LN2). **c** Volcano plot to show fold change in levels of gene expression. **d** Venn diagram showing unique and common expression of genes in parental and resistant cells. **e** Upregulation of Trib2 mRNA in LNCaP-ENR and PCA-2B-ENR cells compared to parental cells by RT-PCR. **f** Western blot showing increased protein levels of Trib2 and targets in enzalutamide-resistant cells, compared to parental cells. **g, h** Representative IHC and quantitation of Trib2 staining intensity in prostate PDX tumors with or without enzalutamide treatment. **i, j** Expression level of Trib2 in patient prostate tumor samples. For panels **h, j** two-tailed unpaired Student’s *t*-test was applied.

To confirm the array data, we detected Trib2 expression by RT-PCR and Western blot which showed strong upregulation of Trib2 mRNA as well as protein levels in enzalutamide-resistant (LNCaP-ENR and PCA-2B-ENR) cells (Fig. 1e, f). Overexpression of Trib2 is correlated with activation of the canonical Akt signaling module showing increased phosphorylation of Akt (pSer-473) and increased protein level of Bcl-xL which are standard markers that promote cell survival and decrease apoptosis. To verify whether the *in vitro* observation of Trib2 overexpression is valid *in vivo*, we analyzed prostate tumors in tissue microarrays and found that prostate PDX tumors overexpress Trib2 when the mice were treated with enzalutamide at 30mg/kg/day for 6 weeks (Fig. 1g, h). Moreover, we found that a vast majority of the clinically advanced metastatic prostate tumors from enzalutamide-treated patients show a robust increase in the expression of Trib2 proteins (Fig. 1i, j). Altogether, these findings suggest that triggering of Trib2 overexpression is a fundamental mechanism in prostate cancer cells both *in vitro* and *in vivo* when treated with enzalutamide, a second-generation direct inhibitor of AR activity.

Prostate cancer cells variably respond to androgenic signaling for proliferation, and the AR-signaling status varies widely among established prostate cancer cell lines. Additionally, some prostate cancer cell lines are devoid of androgen receptor expression and are naturally independent of androgenic stimulus. This provided us with an opportunity to find whether there is any correlation between AR function and the expression levels of Trib2. To address this, we analyzed a range of prostate cancer cell lines, and found that the Trib2 protein level is low in prostate cancer cells with high AR activity, while the level of Trib2 protein is high in prostate cancer cells where the AR signaling function is lowered by drug treatment or is naturally low or lost due to mutation or deletion of the AR gene (Fig. 2a, b). These findings raised a fundamental question whether a functionally negative correlation exists between AR-mediated signaling and the expression of the Trib2 gene.

**Fig. 2.**
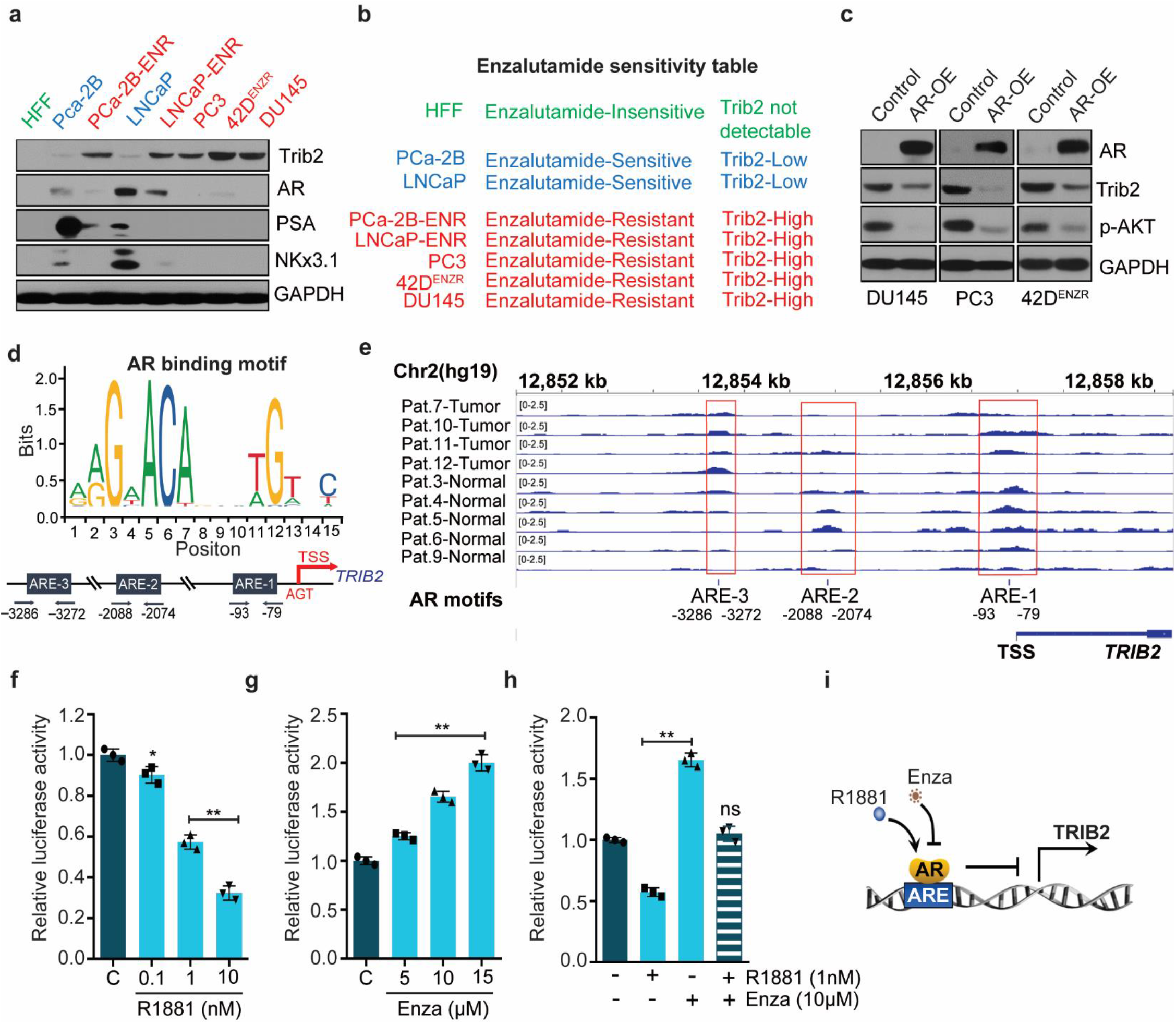
Trib2 is downregulated by AR activity. **a** Whole cell lysates were analyzed by Western blot to detect levels of Trib2, AR and targets. **b** Table showing relationship between Trib2 level and sensitivity to enzalutamide. High level of Trib2 in prostate cancer cells develop enzalutamide-resistance. **c** AR null cells were transfected with full length human AR gene to overexpress (OE) AR proteins. Whole cell lysates were analyzed by Western blot to detect relative expression of AR and Trib2. pAkt is a target of Trib2 which is also downregulated when Trib2 is downregulated by overexpression of AR. **d** Bioinformatics analysis showing AR binding motif obtained from JASPAR database. **e** Genomic representation of AR enrichment on the Trib2 gene in prostate cancer patient samples (GSE70079) mined from ChIP-Seq datasets. **f, g, h** Luciferase reporter activity of Trib2 in LNCaP cells treated either with the synthetic androgen (R1881) or enzalutamide (Enza) for 24 hours. **i** Illustration showing AR signaling mediated transcriptional regulation of Trib2. Data presented as mean values ± SE. \**p* <0.05; \*\**p*<0.005. For panels **f, g, h** One-way ANOVA, Dunnett’s multiple-comparison test was applied.

If Trib2 is negatively regulated by AR signaling, then it is expected that re-introduction and activation of AR in AR-null cells will interfere with Trib2 expression under normal cell culture conditions. Thus, we transfected the high Trib2-expressing, AR-negative/low cells (PC3, DU145, 42D^ENZR^) with full length human AR gene constructs and found that the Trib2 protein level was significantly decreased when the cells were transfected to overexpress AR (Fig. 2c). Thus, reactivation of AR signaling in AR-negative cells (which are naturally high in Trib2 protein levels) inhibits the expression level of Trib2. This finding, together with our previous observation of the upregulation of Trib2 by enzalutamide, suggest that the expression of Trib2 is negatively regulated by AR-signaling.

To address this, we sought to examine the effect of AR transcriptional activity on Trib2 gene expression. We suspected that the promoter of the Trib2 gene may have androgen-response elements (AREs) which presumably bind with AR and in turn inhibit the promoter activity to suppress Trib2 expression. Bioinformatic analysis and ChIP-seq data indicated presence of several AREs within ~5kb promoter region upstream of transcription start site (TSS) of Trib2 gene (Fig. 2d, e), supporting the concept that ligand-bound AR may bind with Trib2 promoter to modulate its expression. To further confirm the AR signaling-mediated transcriptional repression of *TRIB2*, we transfected LNCaP and MDA-PCA-2B cells with Trib2 promoter-luciferase constructs and found that activation of AR by R1881 decreased luciferase activity, while inhibition of AR-signaling by enzalutamide increased the luciferase activity in a concentration-dependent manner (Fig. 2f-h). These findings suggest that the active AR may interact with ARE(s) and in turn inhibits the expression of *TRIB2* via a negative regulation (Fig. 2i), as observed in some other genes, such as BRN2 and SPINK1 (*19,20*).

Because overexpression of Trib2 was observed both in enzalutamide-treated cells and tumors, we asked the question whether Trib2 plays any role in these cells. No specific, target-validated inhibitor of Trib2 is commercially available for molecular and *in vivo* preclinical studies. Thus, we used shRNA-mediated knockdown and found that downregulation of Trib2 strongly inhibits the viability and colony growth of enzalutamide-resistant cells (Fig. 3a-c). Interestingly, normal, non-cancer cells, such as human foreskin fibroblasts (HFF), which do not express detectable Trib2 proteins, remained unaffected with Trib2 shRNA, suggesting that Trib2 plays a critical but selective role in enzalutamide-resistant cells. Recently, it was demonstrated that the EGFR kinase inhibitor, Afatinib (AFA), destabilizes Trib2 protein by covalent modification and primes Trib2 for degradation (*21*). Thus, we also examined the effect of AFA and found that it strongly downregulates Trib2 protein level and kills enzalutamide-resistant cells by triggering apoptosis (Fig. 3e, f). These findings indicate that suitable Trib2-targeting agents could be developed or repurposed as new therapeutic agents to selectively kill and eliminate enzalutamide-resistant prostate cancer cells.

**Fig. 3.**
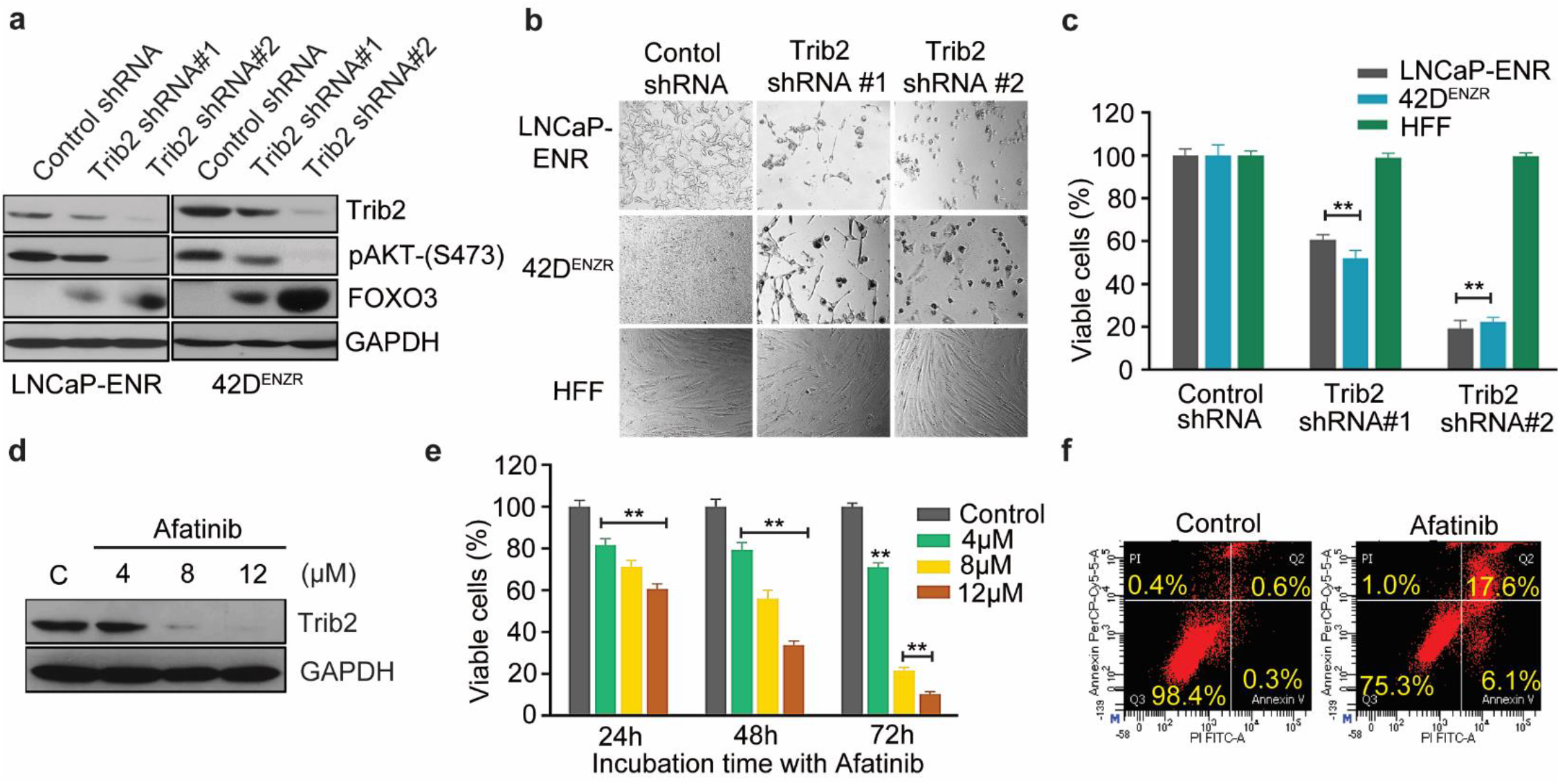
Inhibition of Trib2 kills enzalutamide-resistant prostate cancer cells via induction of apoptosis: **a** Cells were plated overnight and treated with gene specific shRNAs for four days. Protein levels of Trib2, pAkt and forkhead box O3-alpha (FOXO3a) were detected by Western blot. **b, c** Morphological alterations and viability of cells were detected four days after shRNA treatment. **d** LNCaP-ENR cells were treated with Afatinib (AFA) for 24 hours and protein levels were detected by Western blot. **e** Time-dependent decrease in viability of LNCaP-ENR cells was measured by MTS/PES assay (Promega). **f** Apoptosis was measured by Annexin V binding with LNCaP-ENR cells after treatment with AFA for 24 hours. Data presented as mean values ± SE. \*\**p*<0.005. For panels **c, e** Two-way ANOVA, Tukey’s multiple-comparison test was applied.

Since, inhibition of Trib2 strongly inhibits the viability of enzalutamide-resistant prostate cancer cells, Trib2 has emerged as a new molecular target. It also raised the question whether overexpression of Trib2 provides any growth advantage or plays an active role in enzalutamide-resistant cells for their defense against enzalutamide therapy. To address this, we transfected LNCaP and PCA-2B cells with full length human Trib2 gene and observed that the Trib2-overexpressing (Trib2-OE) cells (LNCaP-Trib2, PCA-2B-Trib2) show increased levels of pro-survival proteins, p-Akt and Bcl-xL, and decreased level of Forkhead Box O3 (FOXO3), a tumor suppressor (Fig. 4a). Also, Trib2-OE cells show increased number and size of colonies, suggesting that Trib2 plays a major role in promoting prostate cancer cell growth (Fig. 4b). Interestingly, overexpression of Trib2 alone makes prostate cancer cells completely resistant to therapeutic doses of enzalutamide (Fig. 4c, d). This resistance is abolished, and cells become sensitive to enzalutamide again when Trib2-OE cells were treated with Trib2 shRNA or Afatinib, suggesting that Trib2 greatly contributes to the aggressive growth and resistance to enzalutamide in prostate cancer cells (Fig. 4e).

**Fig. 4.**
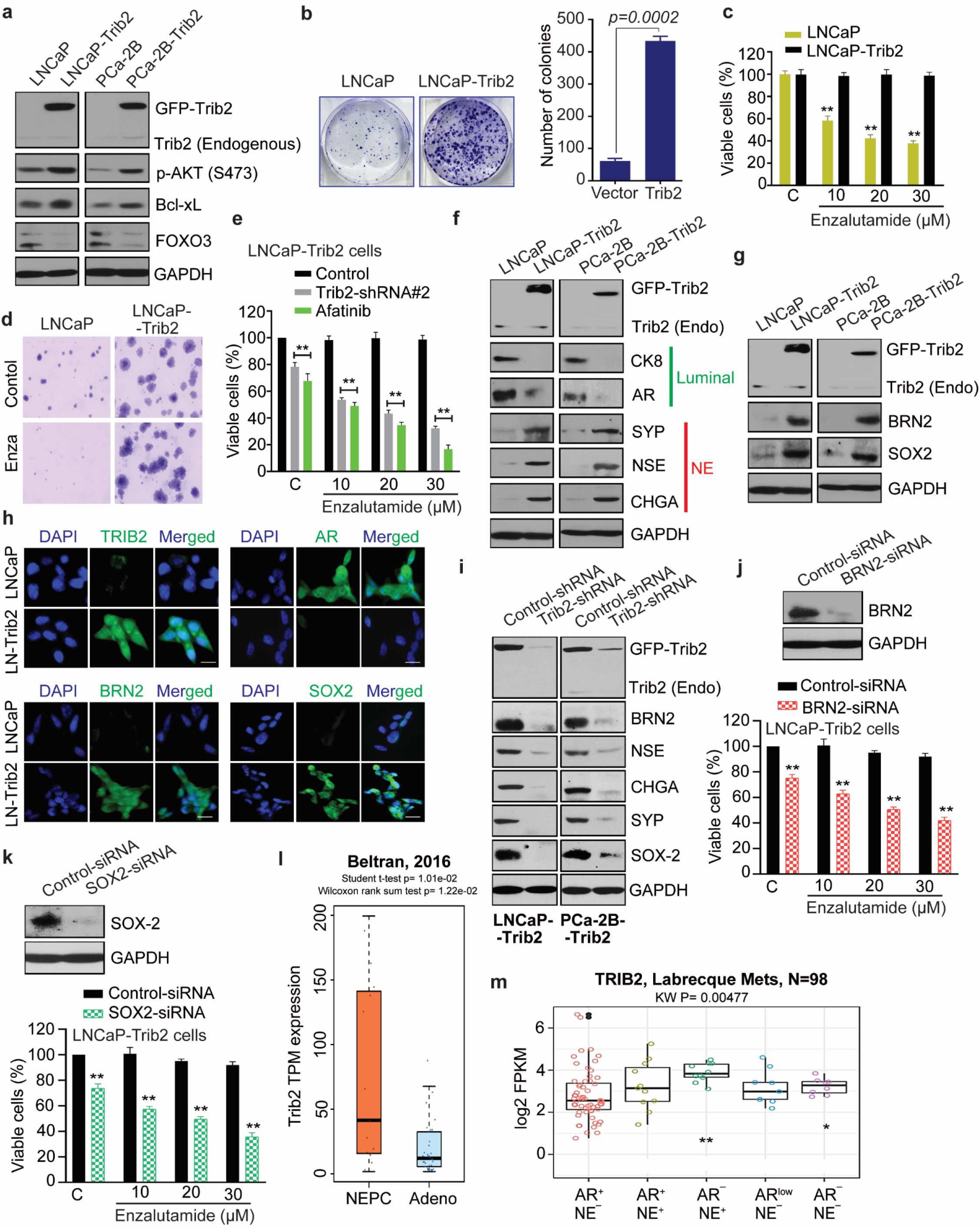
Trib2 enhances prostate cancer cell growth and confers resistance to enzalutamide by promoting lineage plasticity. **a** Immunoblot analysis of indicated protein levels in Trib2-OE cells. **b** Colony growth of transfected and parental cells were analyzed after staining with 0.025% crystal violet. **c** Sensitivity of parental and Trib2-OE cells to enzalutamide was measured by MTS/PES assay. **d** Impact of Trib2 overexpression on enzalutamide resistance was measured by soft-agar colony formation assay. **e** Role of Trib2 in enzalutamide resistance was verified by treating LNCaP-Trib2 cells with shRNA, or by Afatinib. **f** Protein expression of selected lineage markers in Trib2-OE cells was detected by western blot. **g, h** Western blot and immunofluorescence analysis showing protein levels of NE (BRN2) and stemness (SOX2) markers in Trib-OE cells, respectively. *Note:* The data presented in **f** and **g** are generated from the same membrane. Trib2 and GAPDH protein bands shown in **f** and **g** are identical. **i** Immunoblot for NE and stemness markers in Trib2-shRNA treated cells. **j, k** Role of BRN2 and SOX2 in enzalutamide resistance was determined by treating Trib2-OE cells with siRNAs. **l** Boxplot showing Trib2 expression in neuroendocrine prostate cancer (NEPC) versus prostate adenocarcinoma (Adeno) patients (*10*). Wilcoxon rank sum test was used to test the difference between the two groups. **m** Trib2 expression in prostate cancer phenotypes (*11*). AR-high prostate cancer (AR^+^/NE^−^), Amphicrine prostate cancer (AR^+^/NE^+^), Neuroendocrine prostate cancer (AR^−^/NE^+^), AR-low prostate cancer (AR^low^/NE^−^), and Double-negative prostate cancer (AR^−^/NE^−^). Results are expressed as log2 fragments per kilobase of transcript per million mapped reads (FPKM). Data presented as mean values ± SE. \**p* <0.05; \*\**p*<0.005. For panels **h,** scale bar represents 10 μm. For panels **c, e, j, k** Two-way ANOVA, Tukey’s multiple-comparison test was applied.

Resistance to enzalutamide invariably develops and mostly occurs in prostate cancer patients within five years since therapy begins (*5–7*). Enzalutamide-resistant prostate cancer assumes deadly phenotype if an AR-independent mechanism is developed, the spectrum of which is yet to be fully characterized. An AR activity low stemness program as well as neuroendocrine (NE) differentiation were repeatedly observed in enzalutamide-resistant cells and tumors (*22, 23, 8-12*). Recently, the SOX2-mediated lineage plasticity and development of NE phenotype have been demonstrated in PTEN ^(-/-)^: Rb1 ^(-/-)^ double knockout mice (24). Moreover, a role of BRN2, was documented in treatment-induced NE differentiation via upregulation of SOX2 (*19*). To understand the downstream mechanism of Trib2 in enzalutamide resistance, we analyzed the lineage characteristics of Trib2-OE cells.

We found that enzalutamide-resistant Trib2-OE cells (LNCaP-Trib2, PCA-2B-Trib2) show a decrease in the protein level of luminal markers (CK8 and AR) and an increase in neuroendocrine markers such as, SYP, NSE and CHGA (Fig. 4f). These findings indicated that Trib2-induced resistance to enzalutamide does not involve reactivation of AR or maintenance of luminal features, rather Trib2 may be involved in trans-differentiation of luminal epithelial cells to develop NE like characteristics. We also found a strong upregulation of the neuronal transcription factor BRN2, and the stemness factor, SOX2 (Fig. 4g, h). The regulation of neuroendocrine and stemness markers by Trib2 was also confirmed by shRNA knockdown of Trib2 in Trib2-OE cells (Fig. 4i). Moreover, we found that inhibition of BRN2 or SOX2 re-sensitizes Trib2-OE cells to enzalutamide treatment, indicating that the molecular mechanism of Trib2 involves upregulation of BRN2 and SOX2, presumably to increase cellular plasticity (Fig. 4j, k). Our findings suggest that Trib2 helps prostate cancer cells to evade enzalutamide therapy, apparently by switching their characteristics from enzalutamide-sensitive luminal cells to develop NE phenotype (Supplementary Fig. 1). A strong positive correlation of Trib2 was also found with features in NE tumor samples (Fig. 4l, m and Supplementary Fig. 2-4). Thus, Trib2 emerges as a novel driver of NE differentiation and a promising molecular target, characterization of which may reveal a new mechanism underlying development of enzalutamide resistance, and to design a targeted strategy for the management of residual, re-growing prostate cancer cells which evade androgen receptor-targeted therapy.

Based on our findings, it appears that Trib2 induces lineage switching by developing stemness and NE characteristics which supports enzalutamide resistant prostate cancer cells in a way that they are no longer required to depend on AR-mediated signaling. Development of AR(-)/low tumors have now been found to occur in as high as 36% of the patients when treated with enzalutamide and this rate is expected to go up with stronger agents to eliminate androgen receptor function (*25*). Thus, Trib2 emerges as a new driver oncogene to support enzalutamide-resistant prostate cancer cells to continue their growth when treated with clinically achievable doses of enzalutamide via downregulation of AR and development of NE characteristics. Trib2 was originally characterized to play a role in the morphogenesis of *Drosophila* (*26*). Later, Trib2 was characterized to activate Akt and ERK via a still obscure mechanism to support aggressive cancer cell growth and therapeutic resistance (*27–30*). From our unprecedented observations in prostate cancer cells, it became apparent that Trib2 is a *bona fide* driver for survival and enhanced growth in enzalutamide-resistant prostate cancer cells, and that Trib2 plays an important role in enzalutamide resistance by promoting lineage plasticity, so that prostate cancer cells can overcome the loss of support caused by interruption of the androgenic signaling axis.

## Acknowledgments

We thank Dr. Martin E. Gleave of the Vancouver Prostate Center (Vancouver, Canada) for providing CRPC PDX slides and Dr. David Goodrich of the Roswell Park Cancer Institute (Buffalo, NY) for providing the slides from PTEN ^(-/-)^: Rb1 ^(-/-)^ double knockout mice for IHC analysis. This work is supported by the Department of Defense Prostate Cancer Research Program, DOD Award No W81XWH-18-2-0013, W81XWH-18-2-0015, W81XWH-18-2-0016, W81XWH-18-2-0017, W81XWH-18-2-0018 and W81XWH-18-2-0019 PCRP Prostate Cancer Biorepository Network (PCBN).

## Author contributions

- **Conception and design:** J. Ghosh, J. Monga
- **Development of methodology:** J. Ghosh, J. Monga, I. Adrianto
- **Acquisition of data:** J. Ghosh, J. Monga, I. Adrianto
- **Analysis and interpretation of data**: J. Ghosh, J. Monga, I. Adrianto, D. Chitale, J. Alumkal, H. Beltran, A. Zoubeidi
- **Writing, review, and/or revision of the manuscript:** J. Ghosh, J. Monga, I. Adrianto, C. Rogers, S. Gadgeel, H. Beltran, A. Zoubeidi
- **Administrative, technical, or material support:** J. Ghosh, J. Monga, I. Adrianto, C. Rogers, S. Gadgeel, D. Chitale, H. Beltran, J. Alumkal, A. Zoubeidi
- **Supervision:** J. Ghosh, J. Monga
- **Pathology:** J. Ghosh, J. Monga, D. Chitale

## Materials and Methods

### Cell lines and tissue culture

LNCaP, MDA-PCA-2B, PC3, DU145 human prostate cancer cell lines and human fore-skin fibroblasts (HFF) were purchased from American Type Culture Collection (Manassas, VA). The enzalutamide-resistant LNCaP-ENR and MDA-PCA-2B-ENR cells were generated by treating with increasing concentrations of enzalutamide over 3 months. The enzalutamide resistant 42D^ENZR^ cells were kindly provided by Dr. Amina Zoubeidi (Vancouver Prostate Center, Vancouver, Canada). Cells were grown in RPMI medium 1640 or DMEM (Invitrogen, Carlsbad, CA) or HPC1 media (Athena Enzyme, Baltimore, MD). All the media were supplemented with 10% FBS and antibiotics. All enzalutamide-resistant (ENR) cells were cultured in the presence of 30μM enzalutamide.

### RNA sequencing and analysis

Total RNA was isolated from exponentially growing cells using spin columns and reagents in the RNeasy Midi Kit following methods provided by the company (Qiagen, Carlsbad, CA). RNA quality check, cDNA synthesis and sequencing were done at the AGTC core facility (Wayne State University, Detroit, MI) using the Illumina Hi-Seq platform. The analysis of differentially expressed genes between groups was conducted using a negative binomial model as implemented in the EdgeR package in R. A gene with a fold-change > 2 (or < 0.5) and a False Discovery Rate (FDR)-adjusted p-value < 0.05 is considered differentially expressed between two experimental conditions. The heatmap of differentially expressed genes was plotted using the gplots heatmap.2 function in R. Each row was scaled so that it has a mean of zero and a standard deviation of one. The volcano plot showing the differentially expressed gene results was generated using the *EnhancedVolcano package in R*.

### Real-Time quantitative PCR

Total RNA was extracted using RNeasy kit (Qiagen) and 1 μg of total RNA was used for cDNA synthesis using SuperScript III First-Strand kit (Invitrogen) according to the manufacturer’s instructions. PCR reaction mixture was prepared using gene specific TaqMan gene expression assay system (Applied Biosystems). qRT-PCR reactions were performed in triplicate using QuantStudio 6 Flex Real-Time fast PCR System (Applied Biosystems) and 2^-ΔΔCt^ values were used to calculate the relative expression level of the target genes compared to controls. GAPDH was used as a normalization control.

### Western blot

Cells were plated in 60 mm diameter plates and at 60-70% confluency they were treated with inhibitors. After treatment, cells were harvested, washed, and lysed in lysis buffer (50mM HEPES buffer, pH 7.4, 150mM NaCl, 1mM EDTA, 1mM orthovanadate, 10mM sodium pyrophosphate, 10mM sodium fluoride, 1% NP-40, and a cocktail of protease inhibitors). Proteins were separated by 12% SDS-PAGE and transferred to nitrocellulose membranes. Membranes were blocked with 5% nonfat-milk solution and blotted with appropriate primary antibody followed by peroxidase-labeled secondary antibody. Bands were visualized by enhanced chemiluminescence detection kit from Pierce Biotech (Rockford, IL). To be accepted as valid, protein blots were analyzed at least in two independent experiments showing similar results. Antibodies against Trib2, pAKT, Bcl-xL, AR, PSA, Nkx3.1, Sox2 and GAPDH were from Santa Cruz Biotechnology (Santa Cruz, CA). Antibodies against BRN2, Enolase-2 (NSE), Synatophysin (SYP), and Chromogranin A (CHGA) were from Cell Signaling (Danvers, MA). Anti-beta-actin antibody was purchased from Sigma Chemical CO (St. Louis, MO).

### Human prostate cancer specimens and Patient derived xenograft (PDX) tumors

Tissue microarray (TMA) with Enzalutamide-treated prostate cancer specimens were obtained from Prostate Cancer Biorepository Network (PCBN), University of Washington, Seattle, WA, USA. The hormone naïve TMA prostate cancer specimens were obtained from Henry Ford Health System (HFHS), Detroit, Michigan, USA after mandatory approval from the Institutional Review Committee. TMA slides created from PDX models (Untreated and enzalutamide treated) were obtained from Dr. Martin E. Gleave, Vancouver Prostate Centre, Vancouver, Canada. The TMAs were stained for Trib2 using immunohistochemistry (IHC).

### Immunohistochemistry

Slides containing formalin-fixed paraffin-embedded sections were incubated at 60°C for 1h and antigen retrieval was carried out in EnVision FLEX target retrieval solution, Low pH (Agilent Dako, S236984-2) in a PT Link instrument (Agilent Dako, PT200). Slides were washed in 1X TBST wash buffer for 5 min, followed by treatment with Peroxidazed 1 (Biocare Medical, PX968M) for 5 min. Nonspecific background was blocked with Background punisher (Biocare Medical, BP974) with a 10 min treatment. The slides were incubated with appropriate primary antibodies diluted in EnVision FLEX Antibody Diluent (Agilent Dako, K800621-2) overnight at 4°C. Slides were washed and incubated with Mach2 Double Stain 1 or 2 (Biocare Medical, MRCT523/525) for 30 min at room temperature. The Slides were developed using ImmPACT DAB Substrate, Peroxidase (HRP) (Vector Labs, SK-4105) and counterstained with Hematoxylin (Agilent DAKO, K800821-2) for 5 min. Slides were rinsed in distilled water, dried and mounted using EcoMount (Biocare Medical, EM897L). The slides were scanned using Aperio CS2 digital pathology scanner. The staining intensity was quantified by QuPath 0.2.3 (Github) digital image analysis software.

### Plasmids and lentiparticles

Trib2-GFP plasmid was a kind gift from Dr. Wolfgang Link (Faro, Portugal). Mission lentiviral transduction particles for Trib2-shRNA and full-length AR plasmid for overexpression, and the siRNAs targeting BRN2 and SOX2 were purchased from Sigma Aldrich (St. Louis, MO). The Trib2 promoter (−5.6kb) plasmid was custom made through Genecopoeia ((Rockville, MD). For transfection of siRNAs, Lipofectamine RNAiMAX was used in accordance with the manufacturer’s guidelines (Life Technologies).

### Cell viability assay

Cells (~3,000 per well) were plated overnight in 96 well tissue culture plates and treated with drugs or appropriate controls. After 72 hours, cell viability was measured by MTS/PES One Solution Cell Titer Assay following manufacturer’s protocol (Promega Corp, Madison, WI).

### Trib2 promoter luciferase assay

LNCaP cells were transfected with a Trib2 promoter-luciferase construct (5.6kb; Genecopoeia) using Lipofectamine 3000 transfection reagent (Invitrogen). Stable transfected cells were plated in 96-well black/clear bottom plates (Thermo Scientific). Cells were serum starved for 8h and stimulated with R1881 at indicated concentrations for 24 h. For anti-androgen treatment, Trib2 promoter-luciferase expressing cells were treated with enzaluatmide for 24 h. Renilla luciferase activity was measured using Renilla-Glo Luciferase Assay System (Promega, Madison, WI) according to the manufacturer’s protocol.

### Soft-agar colony formation assay

LNCaP and LNCaP-Trib2 cells (1×10^5^) were mixed in 0.5 ml of 0.3% soft-agar and seeded on the top of a 2 ml base layer of 0.6% agar. Plates were allowed to settle and then the agar layers were covered with 2 ml fresh RPMI media containing 10% FBS. Cells were treated with enzalutamide (30μM) for three weeks. Cell growth medium and enzalutamide were exchanged every fourth day. At the end of incubation, cells were stained with 0.025% crystal violet and colonies were counted and photographed under a Leica microscope at x150.

### Statistical analysis

Statistical significance was assessed by either one-way ANOVA, two-way ANOVA with post hoc multiple comparisons test or the two-tailed Student’s *t*-test using GraphPad prism software. The *P*-value of less than 0.05 was defined as significant. Results are expressed as the mean ± standard error of the mean (SEM) from at least three independent experiments and are described in each figure legend when applied.

---

**Supplementary Figure 1.**
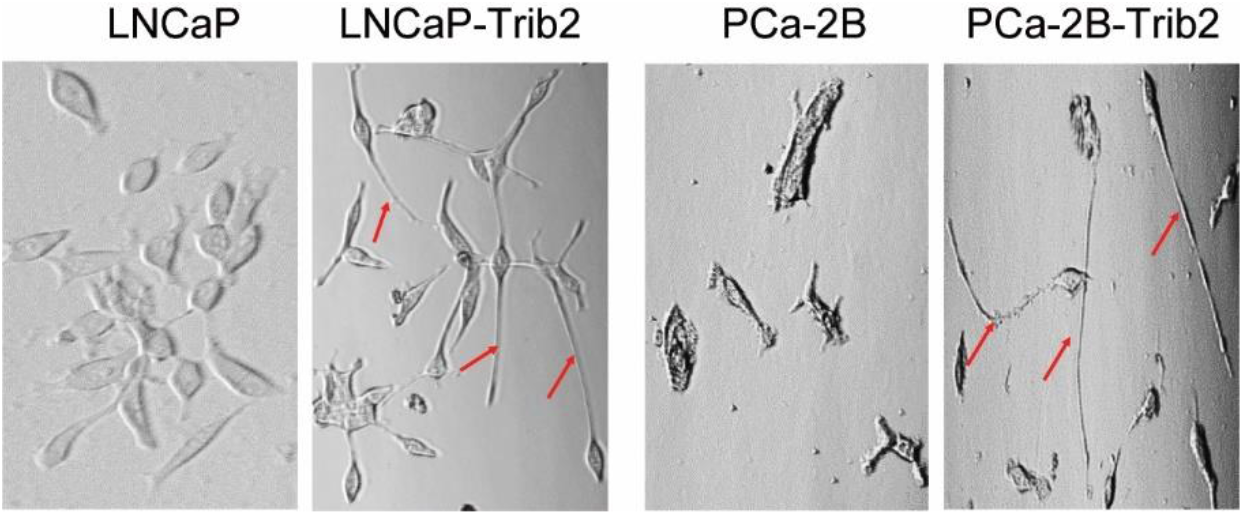
Induction of neuronal morphology. Representative phase-contrast images of Trib2-OE prostate cancer cells. Red arrows indicate extended cellular outgrowth (neurites).

**Supplementary Figure 2.**
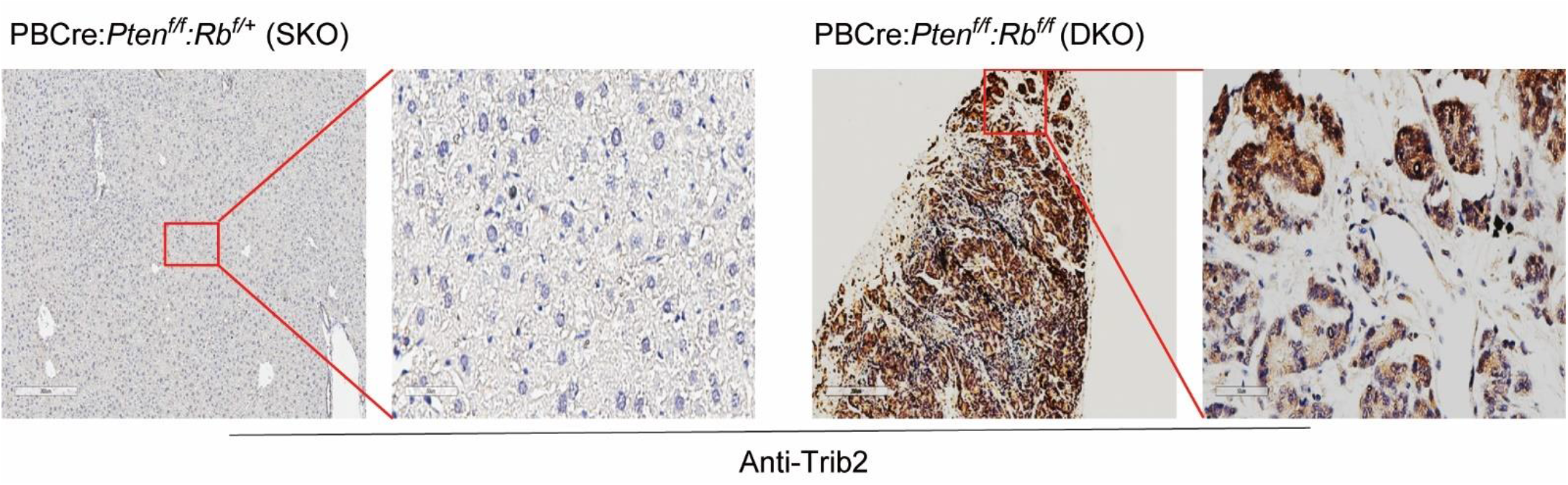
Transgenic mouse model representing neuroendocrine prostate cancer phenotype expresses high levels of Trib2. Tumor sections stained with Trib2 antibody. SKO: Single knockout; DKO: Double knockout.

**Supplementary Figure 3.**
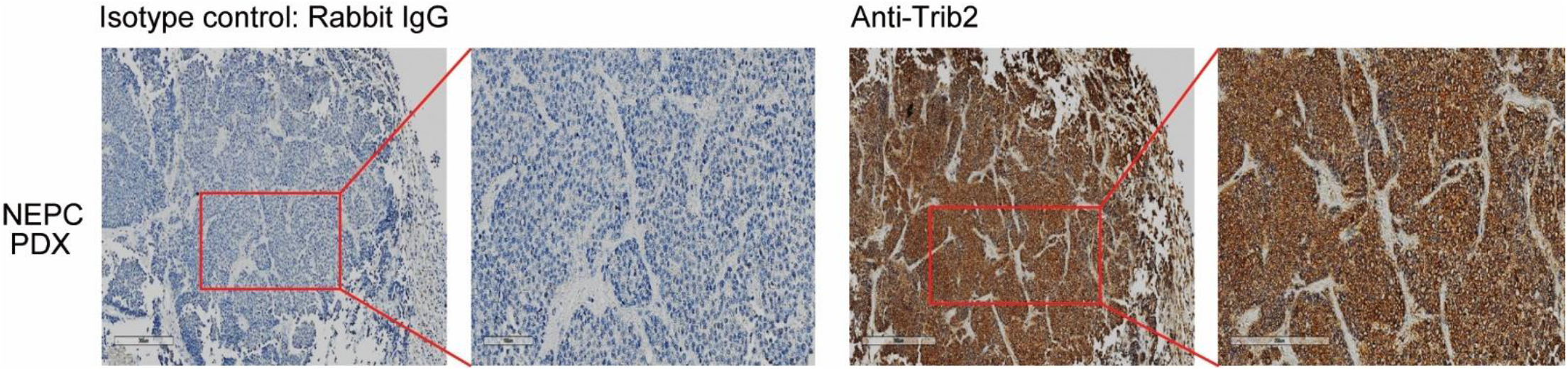
Neuroendocrine prostate cancer patient-derived xenografts (NEPC-PDX) show high levels of Trib2 protein. Representative pictures of IHC analysis of NEPC-PDX tumors (n=3) stained with control IgG or anti-Trib2 antibody.

**Supplementary Figure 4.**
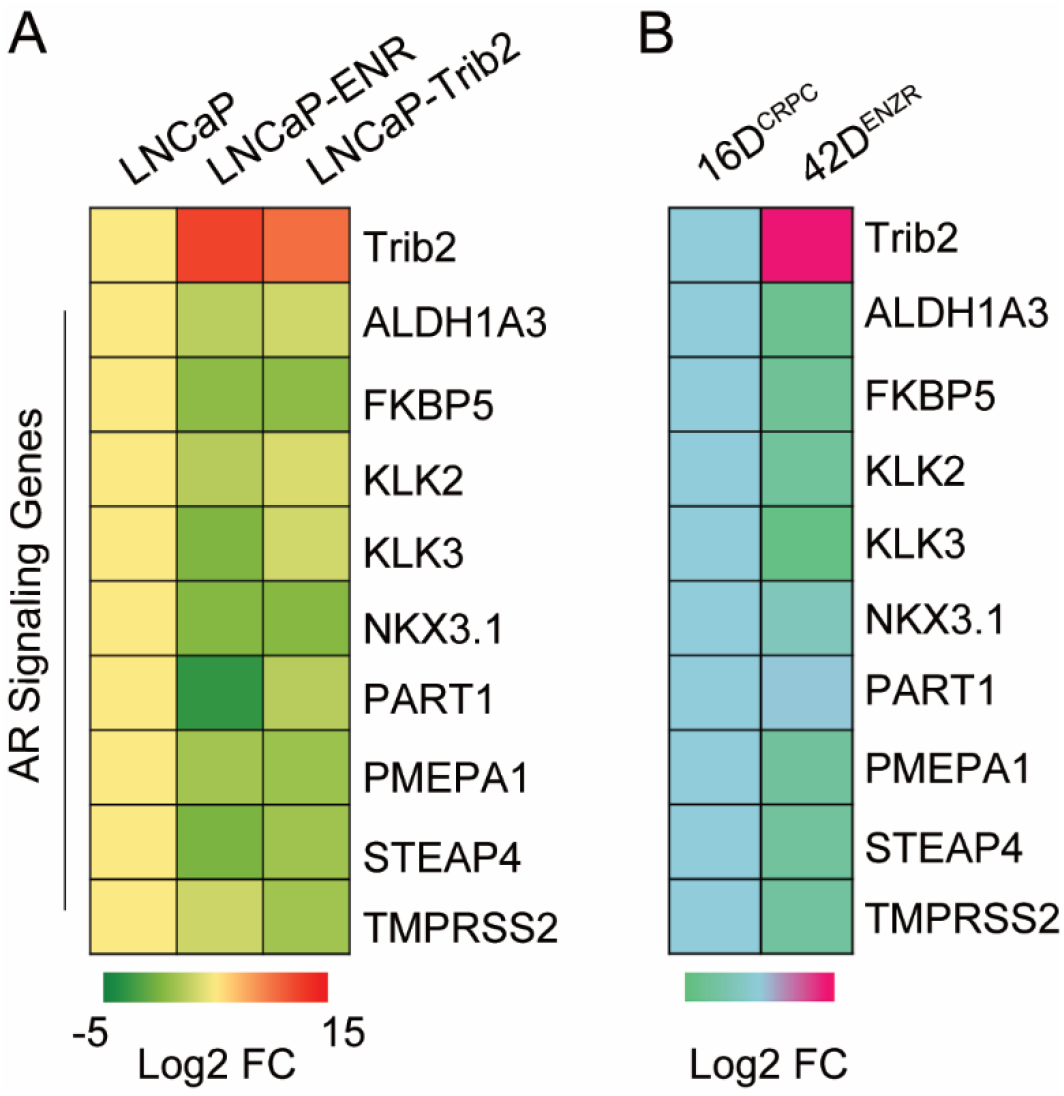
Enzalutamide-resistant prostate cancer cells show high levels of Trib2 expression and low levels of AR signaling markers. Heatmaps showing fold change in Trib2 gene and AR target genes in enzalutamide-resistant prostate cancer cell models compared to parental LNCaP cells.

